# Terroir and rootstock effects on leaf shape in California Central Valley vineyards

**DOI:** 10.1101/2024.04.02.587833

**Authors:** Zoë Migicovsky, Joel F. Swift, Mani Awale, Zachary Helget, Laura L. Klein, Leah Pinkner, Karoline Woodhouse, Peter Cousins, Anne Y. Fennell, Allison J. Miller, Daniel H. Chitwood

**Author notes:** Corresponding authors: Daniel H. Chitwood, Zoë Migicovsky.

## Abstract

**Summary:**

- Embedded in a single leaf shape are the latent signatures of genetic, developmental, and environmental effects. In viticulture, choice of location and rootstock are important decisions that affect the performance and production of the shoot. We hypothesize that these effects influence plant morphology, as reflected in leaf shape.
- We sample 1879 leaves arising from scion and rootstock combinations from commercial vineyards in the Central Valley of California. Our design tests 20 pairwise contrasts between Cabernet Sauvignon and Chardonnay scions from San Joaquin, Merced, and Madera counties from vines grafted to Teleki 5C, 1103 Paulsen, and Freedom rootstocks.
- Using geometric morphometric approaches, we visualize a morphospace in which, in addition to clear separation of Cabernet Sauvignon and Chardonnay scion leaf shapes, an orthogonal source of shape variation affects both varieties. Comparing the Procrustes distances to within and between group means, the additional source of variance is found to arise from location and rootstock effects.
- We describe and visualize a specific shape feature, the angle of the proximal lobe to the midvein that defines the closure of the petiolar sinus, that is attributable to location and rootstock effects and orthogonal to and separate from genetic, developmental, or allometric effects attributable to leaf size.

**Societal Impact Statement (EN):** The innumerable effects of terroir—including climate, soil, microbial environment, biotic interactions, and cultivation practice—collectively alter plant performance and production. A more direct agricultural intervention is grafting, in which genetically distinct shoot and root genotypes are surgically combined to create a chimera that alter shoot performance at a distance. Selection of location and rootstock are intentional decisions in viticulture to positively alter production outcomes. Here, we show that terroir and rootstock alter the shapes of grapevine leaves in commercial vineyards throughout the California Central Valley, documenting the profound effects of these agricultural interventions that alter plant morphology.

## Introduction

Every leaf has only one shape, but that shape is the result of innumerable effects whose signatures are left behind, if only we have the right tools to measure them. Using geometric morphometric methods to quantify shape, these effects can be statistically measured and separated from each other, revealing latent shapes that together comprise leaf morphology (Chitwood and Sinha, 2016). All leaves arising from *Vitis* species are palmate with five lobes, creating a geometric framework in which features are comparable between leaves and species. This framework was leveraged by early ampelographers (after the Greek *ampelos*, άμπελος, literally vine; named after the satyr lover of Dionysus that was the personification of grapevines; Nonnus of Panopolis, *Dionysiaca*, Book 12) to distinguish newly introduced North American rootstock species to 19th century France (Goethe, 1876; 1878; Ravaz, 1902) and eventually wine grape varieties in the 20th century (Galet 1979; 1985; 1988; 1990; 2000). The unique geometrical properties of grapevine leaves led to the application of rigorous mathematical approaches to calculate a mean grapevine leaf while preserving intricate details, like the serrations (Martínez and Grenan, 1999). This mathematical framework is the foundation of geometric morphometric methods, in which statistical sampling of high numbers of leaves can resolve underlying genetic (Chitwood et al., 2014; Klein et al., 2017; Demmings et al., 2019; Chitwood, 2021), developmental (Chitwood et al., 2016a; Bryson et al., 2020), and environmental (Chitwood et al., 2016b; Baumgartner et al., 2020; Chitwood et al., 2021) contributions to grapevine leaf shape.

Yet, even though the field of ampelography was initially created to distinguish shoots of North American rootstock species (Ravaz, 1902), the effects of these roots on grafted scions remains understudied (Migicovsky et al., 2019; Harris et al., 2021). The power of rootstocks lies in the ability for a different genotype than the scion to non-cell autonomously alter the shoot phenotype at a distance (Frank and Chitwood, 2016; Warchefsky et al., 2016; Gaut et al., 2019; Williams et al., 2021). As leaves are a primary constituent of shoot systems, it is reasonable to ask if rootstocks can influence leaf shape. For example, grafting dominant *Me* tomato (*Solanum lycopersicum*) mutant roots to non-mutant shoots results in translocation of the associated mutant KNOTTED1-like homeobox transcript and induces leaf shape changes (Kim et al., 2001).

Here, we apply geometric morphometric approaches to describe the influence of rootstock and location on grapevine leaf shape. We collected 1879 Cabernet Sauvignon and Chardonnay leaves during 2018 and 2019 from commercial vineyards in San Joaquin, Merced, and Madera counties in the Central Valley of California grafted to Teleki 5C, 1103 Paulsen, and Freedom rootstocks. Based on these leaves, we describe a specific shape feature—the angle of the proximal lobe to the midvein that defines the closure of the petiolar sinus—that statistically varies by rootstock and location, and is orthogonal and distinct from genetic, developmental, and allometric effects of leaf size.

## Materials and Methods

### Experimental design

Prior to sampling in 2018, commercial vineyards plots (each with a unique rootstock by scion combination) were selected in San Joaquin, Merced, and Madera county. The study sites include temperate, dry and hot summer (San Joaquin County) and arid and hot steppe (Merced and Madera Counties) climates according to the Köppen-Geiger classification system (**Figure 1A**, Chen and Chen, 2013). Vines with Cabernet Sauvignon and Chardonnay scions on Teleki 5C, 1103 Paulsen, and Freedom rootstocks were sampled during the 2018 and 2019 growing seasons. In San Joaquin County, all scion and rootstock combinations were present, and all scion and location combinations were sampled for the rootstock Freedom. Only select comparisons could be made for rootstock and scion combinations in the Merced and Madera locations and rootstocks. We chose to analyze contrasting pairs of scion, rootstock, and location combinations, where only one rootstock or location contrast is made at a time. 20 such contrasts are present in this study, each identified by number (**Figure 1B**). As described below, this pairwise contrast design aligns with the morphometric methods we use, in which the overall similarity between two shapes is measured as a Procrustes distance.

**Figure 1:**
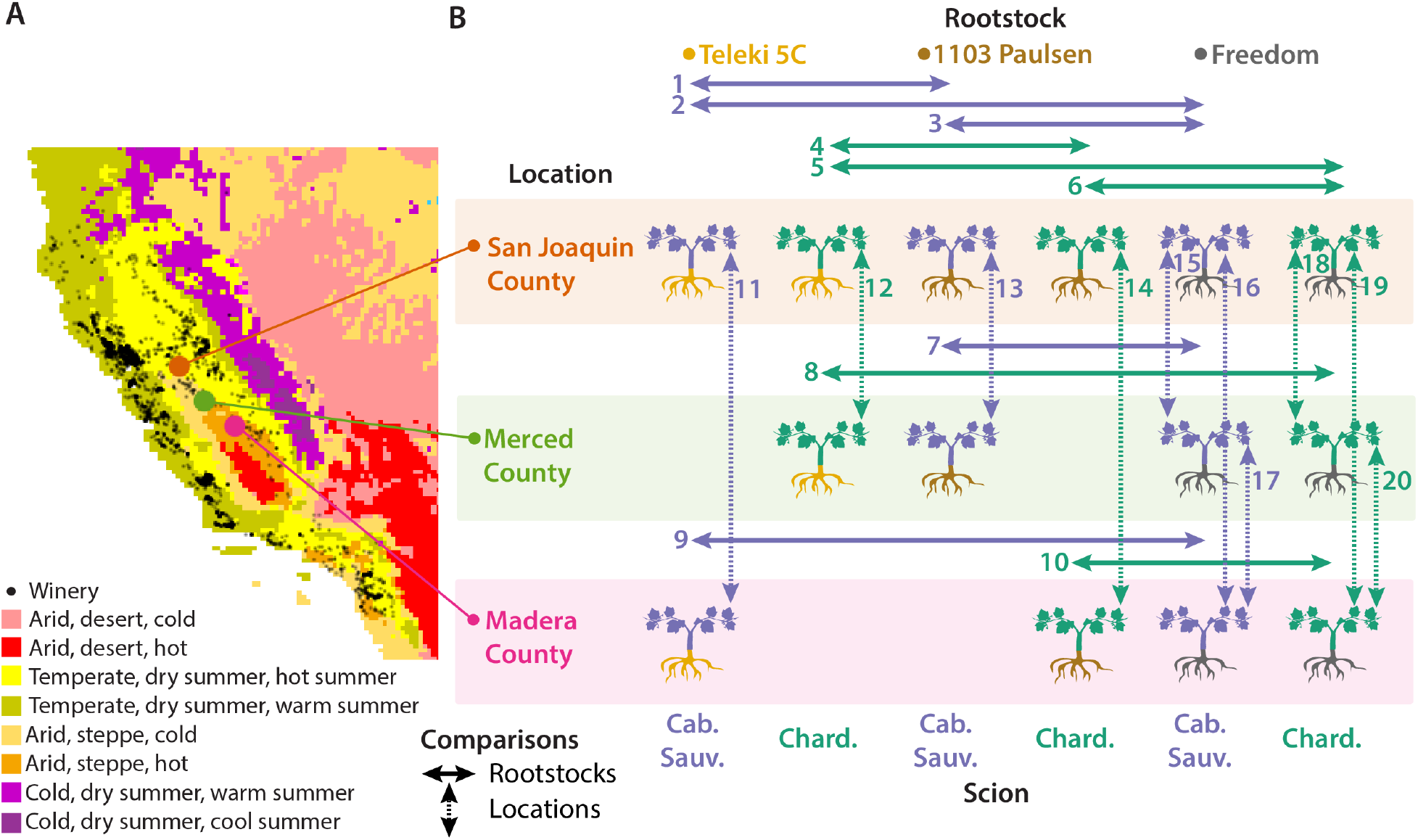
Experimental design. **A)** A map of bonded California winery locations (black points) projected onto Köppen-Geiger climate classifications (see legend). **B)** Sampling design of Cabernet Sauvignon (purple) and Chardonnay (dark green) scions across vineyards in San Joaquin (orange), Merced (light green), and Madera (magenta) counties and Teleki 5C (yellow), 1103 Paulsen (brown), and Freedom (charcoal) rootstocks. 20 contrasts that evaluate effects of pairs of rootstocks (solid, horizontal arrows) or locations (dotted, vertical arrows) are indicated by number.

### Data collection

The vineyard location sampled in San Joaquin is described in detail in Migicovsky et al. (2021; 2023). Due to differing vineyard orientations, leaves were collected from either the north- or west-facing side of the vine. Leaves were sampled from each of the three vineyards weekly in 2018 for 7 weeks from June 19th to August 9th, and for 6 weeks in 2019 from June 11th to July 31st. Vines sampled in 2018 were resampled for the 2019 season, with the exception of those sampled in the last week of 2018 due to the reduction in sampling weeks in 2019. Three vines were sampled for each vineyard block on each sampling date. A LI-6800 Portable Photosynthesis System (LI-Cor Biosciences, Lincoln, NE, USA) was used on two fully expanded mature sunlit leaves on each vine to measure physiological traits between the hours of 10:30am to 2:30pm PST (7:30 to 11:30pm UTC). For LI-6800 measurements, the following parameters were kept constant: flow (600 μmols^-1^), H_2_0 (RH_air 50%), CO_2_ (CO2_r 400 μmol mol^-1^), temperature (Tleaf 33°C), and light (1800 μmol m^-2^ s^-1^). At each timepoint, photosynthetic CO2 rate (A, μmol m^-2^s^-1^) and transpiration rate (E, mol m^-2^s^-1^) were measured. These measurements were also used to calculate instantaneous Water Use Efficiency (WUEi) as A/E (μmol/mol).

A single undamaged shoot was selected from each vine and three leaves were collected from that shoot: the youngest fully expanded leaf, a leaf roughly in the middle of the shoot, and the oldest intact leaf closest to the shoot base. Leaves were trimmed of petioles, placed in a plastic bag on top of one another in order (young, middle, and oldest), and stored in a cooler. Each leaf had its abaxial surface scanned using either a DS-50,000 (Epson, Suwa, Japan) or CanoScan LiDE 220 (Canon, Ōta, Japan) scanner in color with a white background at 1200 dpi. Resulting images were saved as .jpeg files. 21 landmarks were placed onto each leaf scan following the protocol of Chitwood et al. (2021) using the open source software ImageJ (Schneider et al, 2012) or Fiji (Schindelin et al., 2012). Coordinates for each landmark were exported as a CSV file which were merged together with metadata to serve as basis for all subsequent analysis (available here: https://github.com/DanChitwood/terroir_and_rootstock/).

### Morphometric and statistical analysis

The overall similarity between two shapes defined by the same number of points with the same number of dimensions can be measured as a Procrustes distance. Using functions of translation, rotation, scaling, and reflection, the Procrustes distance, calculated between the corresponding points of each shape, is minimized (Goodall, 1991). A population of shapes can be superimposed upon each other through the Generalized Procrustes Analysis (GPA) algorithm (Gower, 1975). Briefly, in GPA an arbitrary shape is chosen as a reference to which all other shapes are aligned. The mean shape is calculated, to which all shapes are again aligned. The algorithm continues until the Procrustes distance between the mean shapes of two cycles falls below an exceedingly low arbitrary threshold. All shapes are then superimposed to the calculated GPA mean shape, allowing corresponding coordinate points to be used comparatively in subsequent statistical analyses.

In this study, 1879 leaf shapes were analyzed. Each leaf consisted of 21 landmarks with 2 coordinates values such that the total dataset size was 1879 x 21 x 2 = 78918. In order to evaluate the overall similarity of two shapes to each other, a Procrustes distance was calculated. To use this measure of similarity to statistically test whether a difference in shape exists between each of the 20 contrasts we evaluated, we used the Kruskal-Wallis one-way analysis of variance to compare within and between group distances to mean leaf shapes. For example, to contrast leaf shapes between groups A and B, we first calculated the GPA means for each group and measured the Procrustes distance of each leaf to its respective mean. We then did the same, but measuring the Procrustes distances of all leaves to the overall common mean. The Kruskal-Wallis one-way analysis of variance was used to determine whether the Procrustes distances of leaves to their respective group means was statistically less than the distance to the overall common mean. A similar method was used to evaluate the physiological data, but instead of using Procrustes distances to a mean shape, the absolute value of residuals to water use efficiency curves modeling photosynthetic rate as a function of transpiration rate were compared between and within groups. We adjusted the reported p values for the contrasts of leaf shape and physiological data using Bonferroni multiple test correction.

All analyses were performed using Python (version 3.10.9) including the numpy (Harris et al., 2020), pandas (McKinney, 2010), matplotlib (Hunter, 2007), and seaborn (Waskom, 2021) modules. The procrustes and stats modules from scipy (Virtanen et al., 2020) were used for Procrustes analysis, Kruskal-Wallis one-way analysis of variance, and the calculation of Spearman correlation coefficients. The PCA function from sklearn (Pedregosa et al., 2011) was used for Principal Component Analysis (PCA) and the inverse transform to calculate eigenleaf shapes of the morphospaces. statsmodels (Seabold and Perktold, 2010) was used for Bonferroni multiple test adjustment. The curve_fit function from scipy.optimize was used to fit curves for water use efficiency (WUE), modeling photosynthetic rate (A, umol m^-2^ s^-1^) as a function of transpiration rate (E, mol m^-2^ s^-1^) using the function A = m*ln(E) - b, where m and b are the estimated slope and intercept of a linear function.

The data and analyses that support the findings of this study are openly available in github at https://github.com/DanChitwood/terroir_and_rootstock.

## Results

In order to visualize the main sources of variance in our data, we performed a Principal Component Analysis (PCA) on the 21 landmarks, each with an x and y coordinate for a total vector length of 42, for all 1879 leaves superimposed using Generalized Procrustes Analysis (GPA) (**Figure 2**). The inverse transform (which given a PCA, provided PC values can return the corresponding leaf shape coordinates) can be used to generate eigenleaves that visualize the variance explained by each PC, or, the morphospace. The first PC explains 62% of the variation in our data and is associated with shape variation corresponding to deeply lobed leaves (low PC1 values) to leaves with less lobing (high PC1 values). Predictably, Cabernet Sauvignon (deeply lobed) and Chardonnay (less lobed) leaves are, with the exception of a few outliers, cleanly separated along this axis and their mean leaf shapes are visibly different from each other, especially with respect to the depth of the sinuses (**Figure 2A**). Despite the clear division of groups based on scion variety, there are a handful of more highly lobed Chardonnay and less lobed Cabernet Sauvignon, as well as several leaves with an intermediate level of lobing that represent a continuum between groups. Given that leaves were sampled at multiple stages of development, and arising from different rootstocks and locations, these seemingly misplaced leaves may represent, for example, a young Cabernet Sauvignon leaf that is less lobed at that stage of development or responding to a unique set of environmental factors. However, shoot position (**Figure 2B**), rootstock (**Figure 2C**) and location (**Figure 2D**), do not correspond in obvious ways to the variance explained by PC1 and PC2 (which combined explain 73% of variance) and the mean leaves of each factor level overlap extensively and can not be differentiated from each other by eye.

**Figure 2:**
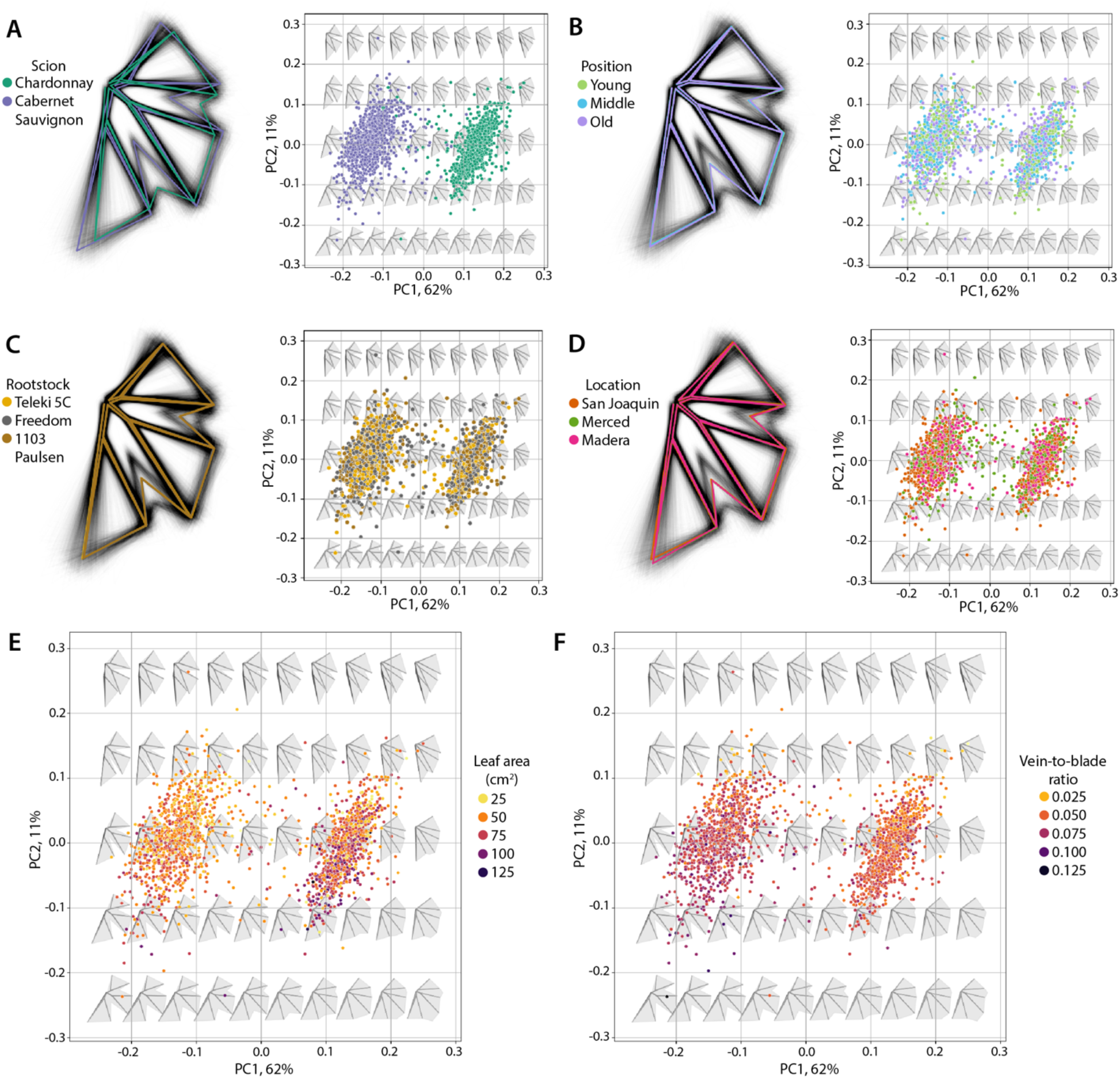
Morphospace. **A)** Principal Component Analysis (PCA) on Generalized Procrustes Analysis (GPA)-adjusted landmarks. Left: Superimposed landmarks of all leaves (black) and the Procrustes mean leaf shapes for Chardonnay (dark green) and Cabernet Sauvignon (purple). Right: eigenleaf representations across the PCA morphospace. Points are colored by scion identity. Similar to (A), panels **(B)**, **(C)**, and **(D)** show superimpositions of Procrustes mean leaves (left) and projections onto the PCA morphospace (right) for shoot position, rootstock, and location factors, respectively (see legends). Allometric indicators of **E)** leaf area (cm^2^) and **F)** vein-to-blade ratio are projected onto the morphospace. Values are indicated by color (see legends).

While PC1 explains over half of the variation in leaf shape, PC2 explains 11% of the variation. Leaves with low PC2 values have proximal lobes with large angles, sometimes exceeding 180°, away from the midvein, covering a larger proportion of the petiolar sinus region, whereas in leaves with high PC2 values the angle between the proximal lobe and midvein is much smaller, sometimes almost 90°, creating a much flatter leaf base in which the proximal lobes on each side of the leaf do not overlap. Both Cabernet Sauvignon and Chardonnay leaves, although separated along PC1, vary along PC2 in similar ways. The similar distributions within each group are suggestive of a shared effect, which we hypothesize may represent leaf size. Previously we demonstrated that the ratio of vein-to-blade area in grapevine leaves is inversely proportional to leaf area (Chitwood et al., 2021) and is a useful indicator of size in normalized leaves arising from morphometric analysis (as is the case here). Projecting leaf area (**Figure 2E**) and vein-to-blade ratio (**Figure 2F**) values on our data, some structure is observed. However, the Spearman correlation coefficients for each of these variables with PC2 is marginal. For Cabernet Sauvignon, the correlation coefficient values for leaf area and vein-to-blade ratio are −0.30 (p value = 5.0 x 10^-21^) and −0.40 (p value = 5.0 x 10^-38^), respectively; for Chardonnay, the correlation coefficient values for leaf area and vein-to-blade ratio are −0.31 (p value = 4.5 x 10^-22^) and −0.25 (p value = 4.2 x 10^-15^), respectively.

To determine if rootstock and location significantly contribute to differences in leaf shape, we turned to the unique structure of our experimental design and the power of using Procrustes distance as a measure of the overall similarity between two leaf shapes. Within our experimental design are 20 contrasts, in which for a pair of scion, rootstock, and locations values, samples differ only by rootstock or by location (**Figure 1**). Reducing our analysis to these 20 contrasts allowed us to leverage the ability of the Procrustes distance metric to compare overall similarity between two samples. By comparing the Procrustes distances of each leaf to the respective mean shape of its group to the distances calculated for each leaf to the overall mean shape using a Kruskal-Wallis one-way analysis of variance, we could assign a p value (multiple test adjusted using Bonferroni) that the leaf shapes of the two contrasting groups differ (**Table 1**).

**Table 1:**
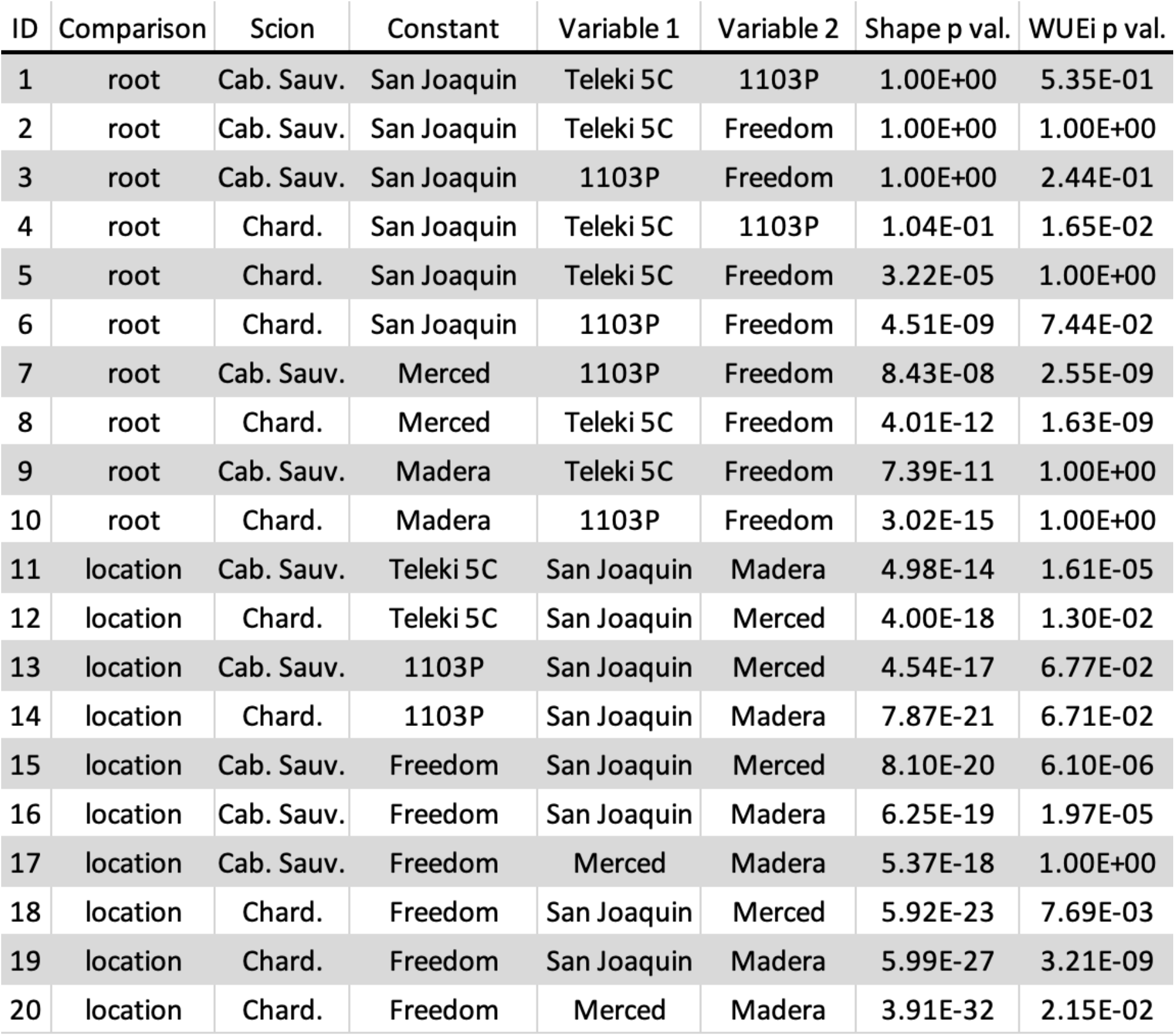
Contrasts by rootstock and location and associated p values for differences in leaf shape and instantaneous water use efficiency (WUEi).

In San Joaquin County, all rootstock contrasts in both Cabernet Sauvignon and Chardonnay were tested, but none of the Cabernet Sauvignon rootstock contrasts were significant. Across all three locations no comparison of Teleki 5C to 1103 Paulsen was significant for either scion. However, all other rootstock and location contrasts were significant. To visualize these shape differences, for each pair of mean leaves, we magnified the difference of each to the other x4 (to see subtle shape effects) and plotted on top of the other mean leaf. Leaves were rotated and scaled so that their midveins overlapped, allowing relative changes in shape to be more easily discerned. There were no qualitative differences in the types of shape differences between rootstock (**Figure 3**) and location (**Figure 4**) contrasts. For some contrasts, slight differences in sinus depth are observed. However, the strongest observable effect was the angle of the proximal lobe to the midvein, similar to the shape variance associated with PC2 described above (**Figure 2**). Consistent directionality (for example, a particular rootstock or location having a wider proximal lobe angle than another) was not immediately obvious.

**Figure 3:**
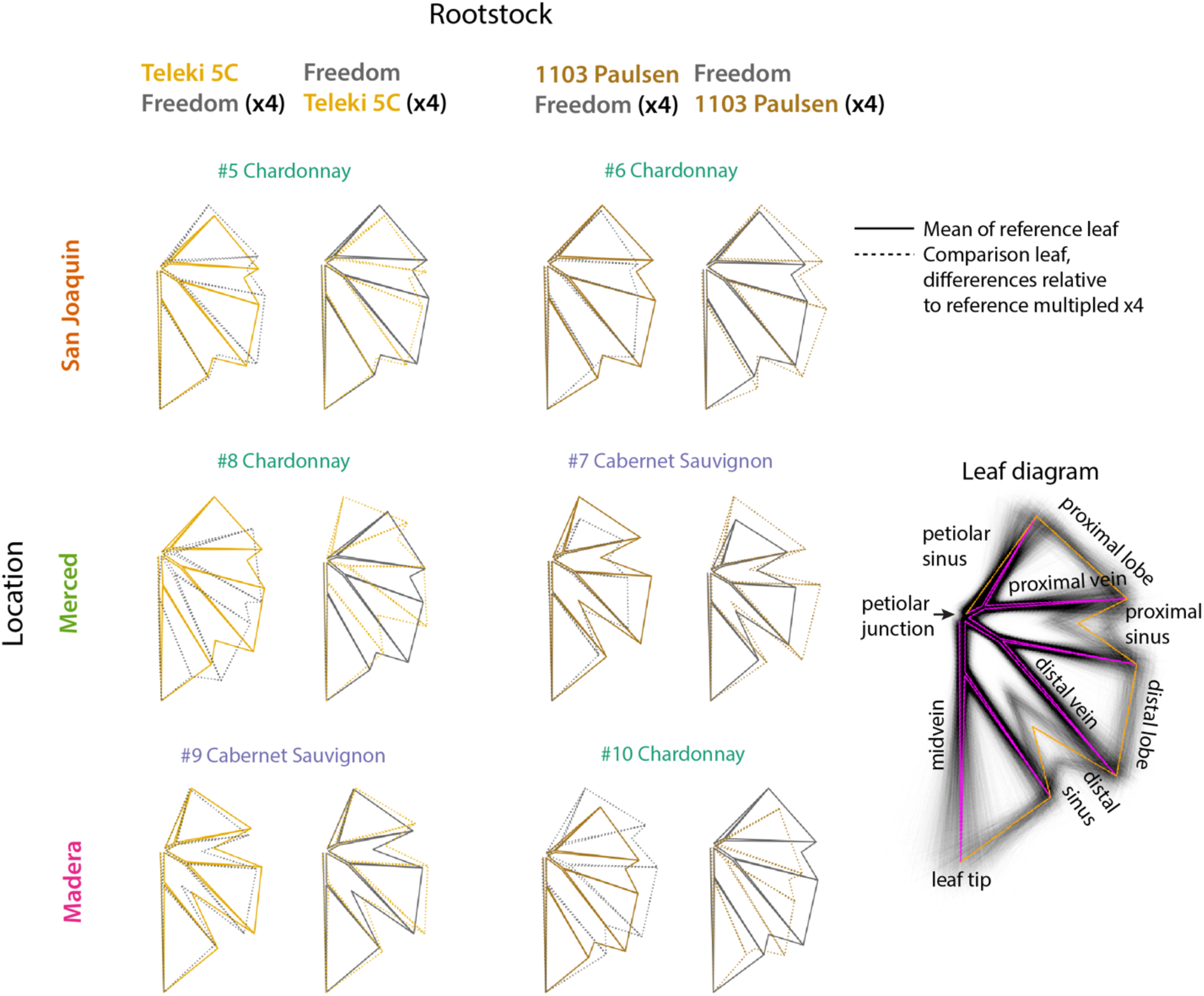
Comparisons of rootstock effects. For each significant rootstock comparison, visualizations of differences between Procrustes mean leaf shapes are visualized as a reference leaf (solid outline) to a comparison leaf (dotted outline), in which the difference to the reference has been multiplied by x4. The differences of each rootstock to the other are visualized in turn. Rootstock pairs are arranged by column and the locations the samples arise from by row. The identification number of each contrast and the scion that was sampled are indicated. A leaf diagram labels morphological features on top of GPA-adjusted leaf outlines and the overall Procrustes mean leaf (orange and magenta).

**Figure 4:**
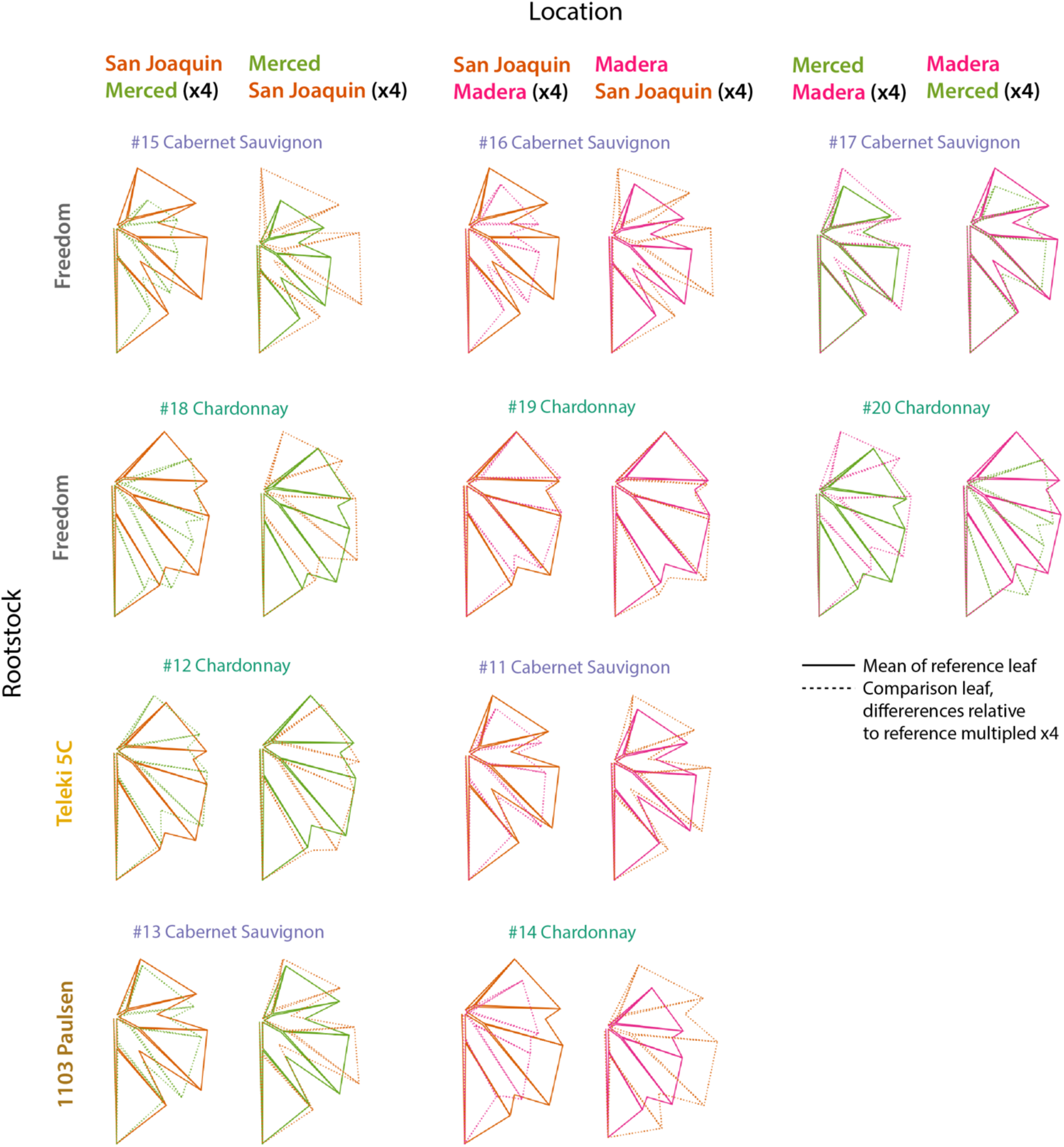
Comparisons of location effects. For each significant location comparison, visualizations of differences between Procrustes mean leaf shapes are visualized as a reference leaf (solid outline) to a comparison leaf (dotted outline), in which the difference to the reference has been multiplied by x4. The differences of each location to the other are visualized in turn. Location pairs are arranged by column and the rootstocks the samples arise from by row. The identification number of each contrast and the scion that was sampled are indicated.

We analyzed the physiological data, collected from the same vines as the leaf shape data, in the same manner as our shape data, so that the results could be directly compared. We first modeled water use efficiency by fitting curves of photosynthetic rate (A, umol m^-2^ s^-1^) versus transpiration rate (E, mol m^-2^ s^-1^) (**Figure 5A**). Differences in the trajectories of the Cabernet Sauvignon and Chardonnay water use efficiency curves can be seen, with Cabernet Sauvignon assimilating at higher rates for a given transpiration rate than Chardonnay (**Figure 5B**), but differences between rootstocks (**Figure 5C**) and locations (**Figure 5D**) are more subtle. Similar to leaf shape, to see if we could detect differences in any of the 20 contrasts, we compared the absolute value of residuals to the fitted curve of samples to their respective group to the absolute value of residuals of the overall fitted curve using a Kruskal-Wallis one-way analysis of variance. After multiple test adjustment, none of the 20 contrasts was statistically significant (data not shown). Instead, for each sample we calculated instantaneous water use efficiency (WUEi, A/E) and tested if any of the 20 contrasts were statistically significant, again using the Kruskal-Wallis test. A number of the contrasts were statistically significant after multiple test adjustment, but there was no obvious overlap in the contrasts that were significant between leaf shape and WUEi (**Table 1**).

**Figure 5:**
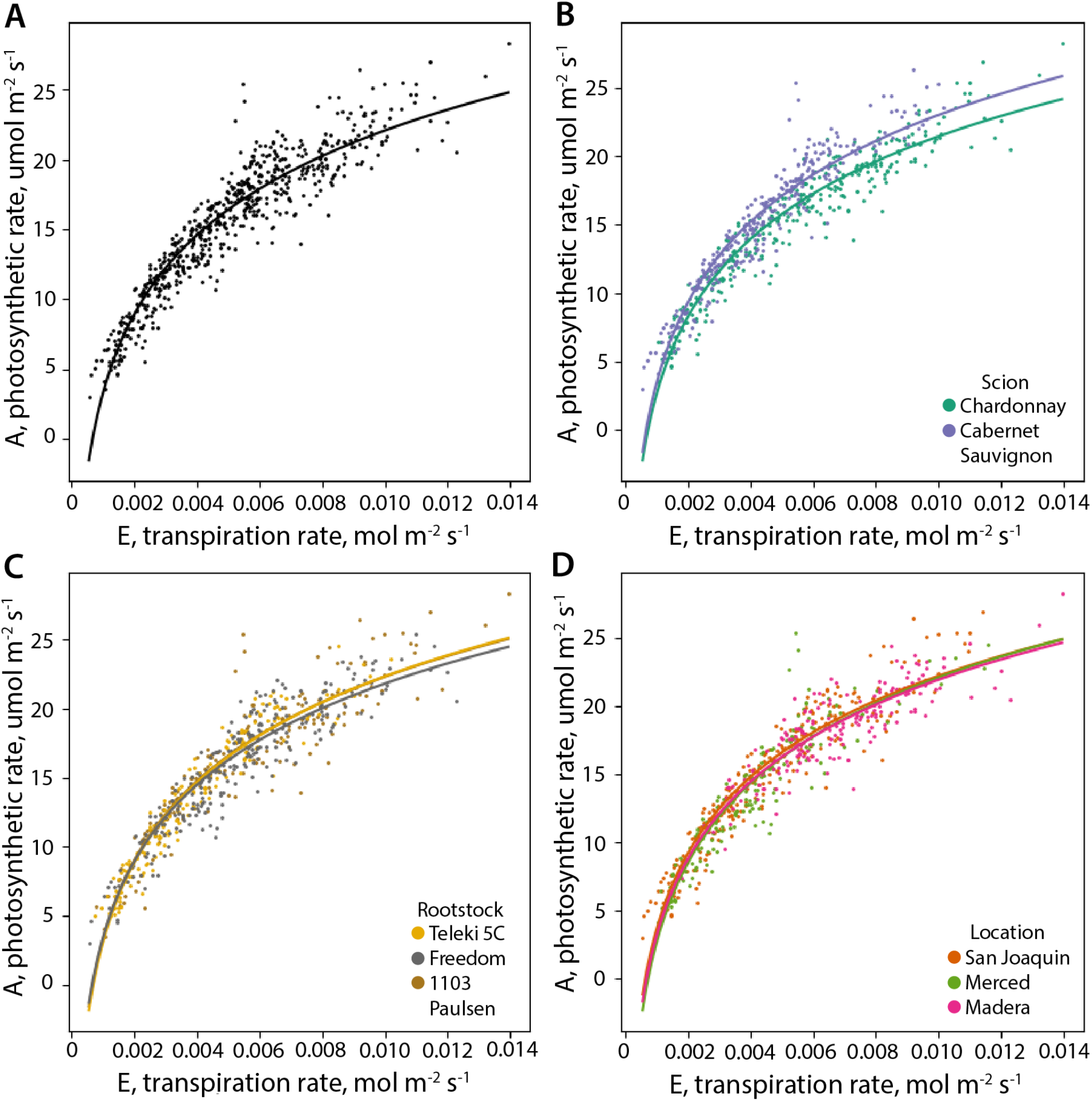
Water use efficiency (WUE) models. **A)** For all samples, photosynthetic rate (A, umol m^-2^ s^-1^) plotted against transpiration rate (E, mol m^-2^ s^-1^). A fitted curve modeling photosynthetic rate as a function of transpiration rate, A = m*ln(E) – b, is shown. Similar to (A), panels **(B)**, **(C)**, and **(D)** show plots and fitted curves by scion, rootstock, and location factors, respectively (see legends).

Our results demonstrate that there are statistical differences in leaf shape between rootstock and location (**Table 1**) and that this shape variation has qualitative similarities, such as variability in the angle of the proximal lobe to the midvein (**Figures 3 and 4**). To isolate these shape differences more specifically, we created morphospaces for Cabernet Sauvignon (**Figure 6A**) and Chardonnay (**Figure 6B**) leaves using only differences in leaf shape between the mean leaves for each significant contrast (magnified x4, as shown in **Figures 3 and 4**). The leaf shapes representing each contrast are connected by line segments, and predictably fall on opposite sides of their respective PC1 axes. The overwhelming observed shape variance along each PC1 axis is the angle of the proximal lobe to the midvein, with nearly 180° angles observed in both Cabernet Sauvignon and Chardonnay for the lowest PC1 values that decrease with higher PC1 values. Directionality is not observed, except for the case of location contrasts in Cabernet Sauvignon leaves, in which leaves from San Joaquin County have larger proximal lobe angles than those from Merced or Madera counties.

**Figure 6:**
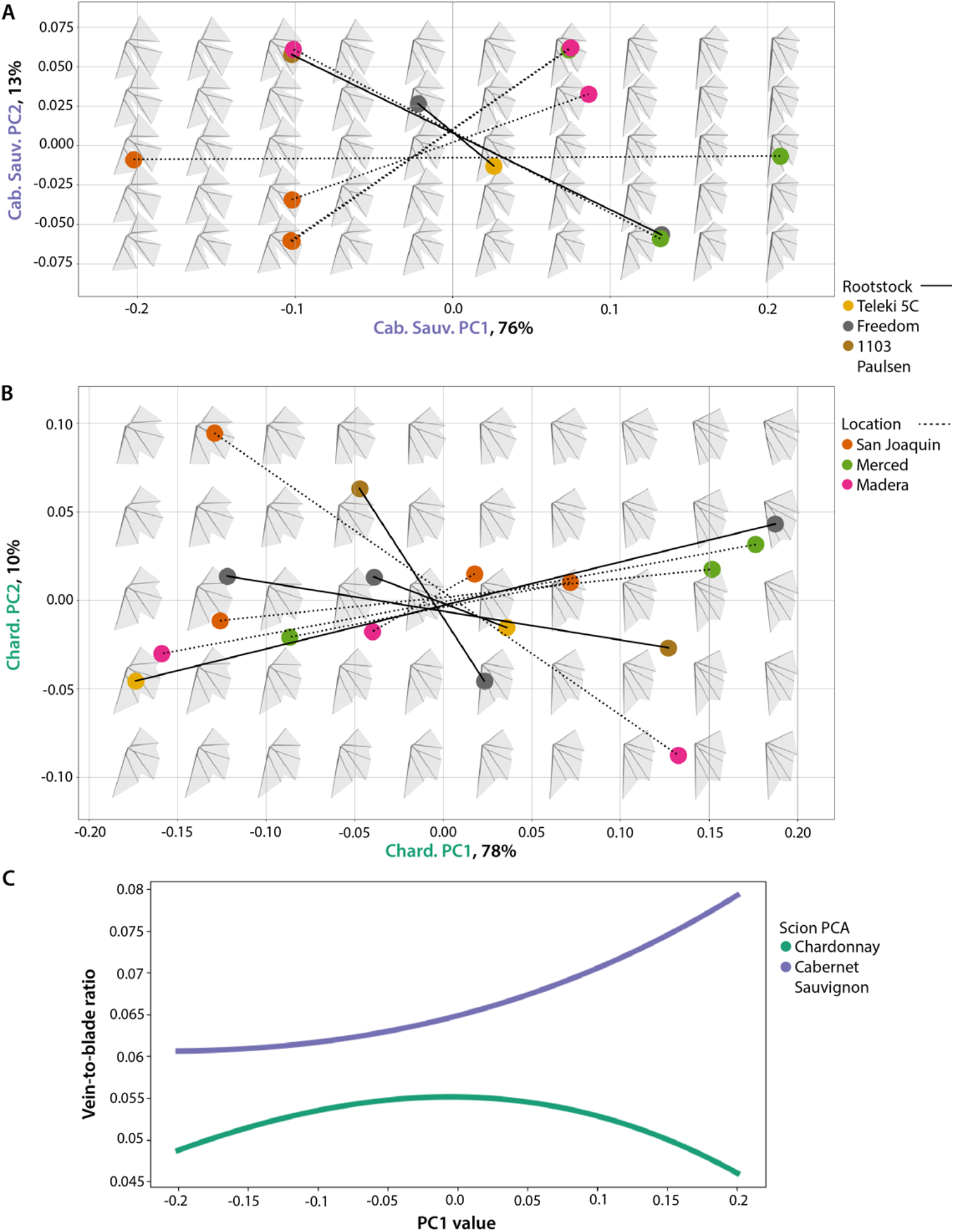
Morphospace of rootstock and location effects on leaf shape. For each significant contrast, divided by Cabernet Sauvignon **(A)** and Chardonnay **(B)** scions, the magnified differences (x4) in leaf shape were used to construct a morphospace. Each pair of contrasted leaf shapes is connected by a line segment indicating the type of comparison, either rootstock (solid) or location (dotted). Points are colored by identity (see legends). Eigenleaf representations are provided to visualize the morphospace. **C)** The modeled vein-to-blade ratio values for eigenleaves across PC1 values for the Cabernet Sauvignon (purple) and Chardonnay (green) PCA morphospaces.

Variation in proximal lobe angle in the PCAs representing individual contrasts by scion (**Figure 6**) reflect the variation observed along PC2 in the overall PCA (**Figure 2**). To determine if this represented an allometric effect related to leaf area, we modeled vein-to-blade ratio (as a proxy of leaf size for normalized leaves) as a function of each PC1 value from **Figure 6A** or **Figure 6B** using eigenleaf representations from the inverse transform (**Figure 6C**). Cabernet Sauvignon values are higher than Chardonnay as expected for a deeply lobed leaf (Migicovsky et al., 2022) and show a marginally positive relationship between PC1 values and vein-to-blade ratio, but Chardonnay does not. The range of vein-to-blade ratios across PC1 values of the PCA of leaf differences is 0.045 to 0.080 (**Figure 6C**), only a fraction of the vein-to-blade ratios observed for actual leaves (0.025 to 0.125, **Figure 2F**). We therefore do not attribute allometric (leaf size) variation to that explaining leaf shape differences by rootstock or location effects.

## Discussion

By measuring leaf shape across three vineyards, two scions, and three rootstocks for a total of 13 weeks across two years, we are able to describe the impact of both terroir and rootstock on altering the shapes of grapevine leaves throughout the California Central Valley. At one location in our study, San Joaquin County, all contrasts for the three rootstocks grafted to both Cabernet Sauvignon and Chardonnay are present. The location was a production vineyard, in which 15 rootstocks are grafted to both scions in a randomized block design. At this location, we have analyzed historical data showing that rootstock choice can modulate yield and vegetative biomass, and even more strongly Ravaz index (the ratio of yield to vegetative biomass) by almost up to 100% (Migicovsky et al., 2021). At the end of 30 years of production at this site, we analyzed the dendrochronology of scion trunk segments using X-ray Computed Tomography, showing that underlying the effects on Ravaz index, rootstocks had altered secondary patterning of the vasculature, and likely the hydraulic performance, of the scion continuously over the life of the vineyard (Migicovsky et al., 2023). Both the effects of rootstocks on Ravaz index and secondary patterning are strongly additive and robust: that is, regardless of scion properties or environmental effects, rootstocks consistently add or subtract from scion trait values. Although we only measured 3 of the 15 rootstocks present in San Joaquin County that are comparable with the other sites, the lack of directionality in rootstock-induced changes in leaf shape (**Figures 3 and 6**) and associated changes in photosynthetic assimilation or transpiration measured coincident with collecting leaves (**Figure 5**, **Table 1**), suggests that leaf shape is not a mechanism affecting plant physiology. Nonetheless, we detect clear changes in leaf shape, orthogonal to genetic differences that define varieties, that arise from effects of rootstocks and location (**Figures 2, 3, 4, and 6**). It is possible that such shape changes represent a constraint that results from changes in vascular patterning, hydraulic flux, the canopy, or the ratio of reproductive to vegetative biomass to which it is grafted.

Although the effects of rootstock and location on leaf shape we are proposing here are new, we previously proposed a framework in which genetic (Chitwood et al., 2014; Klein et al., 2017; Demmings et al., 2019; Chitwood, 2021), developmental (Chitwood et al., 2016a; Bryson et al., 2020), and year-to-year variation in responses to the environment (Chitwood et al., 2016b; Chitwood et al., 2021) are orthogonal and separate to each other (Chitwood and Topp, 2015; Chitwood and Mullins, 2022). Leaf shape variation associated with rootstock and location is strikingly orthogonal to the changes in leaf shape that strongly separate Cabernet Sauvignon and Chardonnay, to the point that it creates very similar distributions in each variety (**Figure 2**). We previously described allometric changes in grapevine leaf shape that are inversely proportional to leaf size across the *Vitis* morphospace, such that the natural log of the ratio of vein area to blade area in a leaf linearly decreases relative to the natural log of leaf area (Chitwood et al., 2021; Chitwood and Mullins, 2022). We reject the hypothesis that the variation orthogonal to the shape differences that define Cabernet Sauvignon and Chardonnay is a random effect due to stochastic sampling of leaf area, since the differences in mean leaf shape, specifically the variation in vein-to-blade ratio (and therefore leaf size) (**Figure 6**), was only a small fraction of the total in the dataset (**Figure 1**). Rather, we believe that the angle of the proximal lobe to the midvein, that defines the petiolar sinus, is a unique leaf morphology trait that varies by rootstock and location arising from physiological effects.

## Conclusion

We describe an additive effect on leaf shape that varies with rootstock and location in grapevine leaves. The angle of the proximal lobe to the midvein modulates the shape of the petiolar sinus. This shape variation is orthogonal to and qualitatively distinct from variation arising from genetic differences across *Vitis* or allometric variation arising from developmental or year-to-year effects that alter leaf size. The variation does not seem to causally affect leaf photosynthetic rates or transpiration, but likely arises as a developmental constraint from changes in overall plant physiology impacted by terroir and rootstock choice.

## Acknowledgements

All authors were supported by the National Science Foundation Plant Genome Research Program award number 1546869. DHC was additionally supported by National Science Foundation Plant Genome Research Program award numbers IOS-2310355, IOS-2310356, and IOS-2310357.

We acknowledge Julie Curless (Missouri State University), Mya Ly(Missouri State University), Vy Nguyen (Missouri State University), Dalton Gilig (University of Missouri), and Ilona Natsch (Saint Louis University) for assistance in sampling and landmarking of the leaves. We would also like to acknowledge Laszlo Kovacs (Missouri State University) and Misha Kwasniewski (The Pennsylvania State University) for student supervisory support.

We thank E & J Gallo Winery and vineyard managers and workers for their generous contributions of time providing safe access and training to sample commercial sites and collect data.

## Author Contributions

Data collection: ZM, JFS, MA, ZH, LLK, LP, KW

Student advising and supervision: ZM, JFS, PC, AYF, AJM, DHC

Conceptualization: ZM, JFS, PC, AYF, AJM, DHC

Data analysis: ZM, JFS, DHC

Writing: ZM, JFS, DHC

Reading and revising: ZM, JFS, MA, ZH, LLK, LP, KW, PC, AYF, AJM, DHC

## Data Availability Statement

The data that support the findings of this study are openly available in github at https://github.com/DanChitwood/terroir_and_rootstock.

